# Circulating soluble urokinase-type plasminogen activator receptor reflects disease severity in a mouse model of diabetic kidney disease and heart failure with preserved ejection fraction

**DOI:** 10.64898/2026.06.30.735488

**Authors:** Sarah Torp Yttergren, Linn Salto Mamsen, Maria Ougaard, Louise Thisted, Henrik H Hansen, Urmas Roostalu

## Abstract

Circulating biomarkers are increasingly used for patient risk stratification in chronic kidney disease (CKD) and heart failure with preserved ejection fraction (HFpEF). However, clinically relevant circulating biomarkers remain insufficiently characterized in rodent models recapitulating diabetic cardiorenal disease with HFpEF. To address this gap, we evaluated 20 translationally relevant inflammation-associated biomarkers in the diabetic db/db uninephrectomized (UNx)-ReninAAV mouse model of CKD and HFpEF. db/db UNx-ReninAAV mice exhibited marked increases in circulating soluble urokinase-type plasminogen activator receptor (suPAR) and monocyte chemoattractant protein-1 (MCP-1), and in interleukin 10 (IL-10) at late stages of disease. Histological analyses confirmed increased tissue expression of suPAR in the heart and kidney and of MCP-1 in the heart. Notably, circulating suPAR levels correlated with disease severity, including systolic and diastolic cardiac dysfunction and albuminuria. Together, these results provide a systematic analysis of biomarkers in a rodent model of diabetes, CKD and HFpEF and identify suPAR as the biomarker most closely associated with disease severity.

## Introduction

Systemic inflammation is associated with and contributes to the progression of heart failure with preserved ejection fraction (HFpEF) and chronic kidney disease (CKD)(1–5). The coexistence of obesity (6–8), type 2 diabetes and hypertension (9,10) further amplifies inflammatory signaling, promoting pathological remodeling in both the heart and the kidney. Consequently, circulating inflammatory cytokines and other associated signaling molecules have emerged as attractive biomarkers for assessing disease severity and treatment response.

A large number of circulating biomarkers have been evaluated for early diagnosis, monitoring and risk stratification in heart failure and other chronic diseases, with inflammation- and fibrosis-associated markers among the most promising (11,12). Among these, soluble urokinase-type plasminogen activator receptor (suPAR), a marker of chronic inflammation, has emerged as a promising prognostic biomarker. Elevated plasma suPAR is associated with progression of diastolic dysfunction (13) and adverse cardiovascular outcomes in HFpEF (14,15). High suPAR level is likewise associated with poor clinical outcomes across a range of cardiovascular, renal and metabolic diseases, including coronary artery disease, heart failure with reduced ejection fraction (HFrEF), diabetes, CKD and cancer (16–20). In the general population, elevated suPAR is associated with incident heart failure, development of CKD, cancer, type 2 diabetes, and all-cause mortality (19,21,22). A number of other inflammation biomarkers, including monocyte chemoattractant protein 1 (MCP1), tumor necrosis factor alpha (TNFα), interleukin-1 (IL-1), IL-6 and IL-8 have been reported in patients with HFpEF (23,24). Despite their clinical relevance, circulating inflammatory biomarkers have not been systematically evaluated in rodent models of HFpEF and diabetic kidney disease. Preclinical studies have largely relied on conventional serum measures of hyperglycemia, renal injury and, in some cases, cardiac damage (e.g. NT-proBNP), whereas systematic characterization of clinically relevant circulating biomarkers remains limited (25–27). Identification of relevant circulating biomarkers would facilitate noninvasive assessment of disease severity and therapeutic efficacy in preclinical studies and improve the translational value of rodent models.

To address this gap, we systematically evaluated the plasma levels of 20 inflammatory biomarkers in db/db UNx-ReninAAV mice, a translational model of diabetes-associated CKD and HFpEF (28–32) and identified suPAR as the biomarker most closely associated with cardiac and renal dysfunction.

## Materials and methods

### Ethics statement

The Danish Animal Experiments Inspectorate approved all experiments, which were conducted using internationally accepted principles for the use of laboratory animals (License No. 2023-15-0201-01456).

### Experimental design

#### Animal model

Female *db/m* and *db/db* (*BKS.CgDock7m+/+ Leprdb/J*) mice (5 weeks old at arrival; 12 weeks at study start) were obtained from Charles River (Calco, Italy). All animals were acclimatized for one week and housed under controlled conditions (12:12-h light-dark cycle; 24 °C, 50 ± 10% humidity). Mice were identified with subcutaneous microchips (PetID Microchip, E-vet, Haderslev, Denmark) and had ad libitum access to tap water and chow (Altromin 1324, Brogaarden, Hørsholm, Denmark).

Temporal progression of plasma inflammatory biomarkers and their association with left ventricular function, cardiac hypertrophy, albuminuria, obesity, and diabetes were investigated in the *db/db* uninephrectomized (UNx)-ReninAAV mouse model. Female *db/db* mice (6 weeks old) received ReninAAV (2.5×10¹⁰ GC/mouse, IV) at week -5, followed by UNx at week -4. Mice were randomized into four groups (n≈12/group) in a running fashion based on body weight and blood glucose, with echocardiography performed (see (32)) at either day 1, weeks 4, 8 or 12, and termination 1–2 days later. Age-matched *db/m* healthy controls (n=13) underwent echocardiography and termination at week 12 together with the final group of *db/db* UNx-ReninAAV mice. At termination, heart and kidney weights were measured, and urine was collected for kidney function analysis (data presented elsewhere (32)). Plasma samples for analysis of biomarkers of inflammation were collected at each termination. Heart and kidney samples for histological assessment were collected from a separate cohort of n=10 *db/db* UNx-ReninAAV and n=9 lean *db/m* healthy control mice. Blood samples and urine were collected for analyses of glycated hemoglobin A1c (HbA1c) and urine albumin to creatinine ratio (uACR), respectively, confirming diabetes, renal failure and cardiac hypertrophy in the second cohort (Supplemental fig 1).

### Plasma and urine biomarker sampling and analysis

#### Spot urine

Spot urine samples were obtained 1–2 days prior to study termination by collecting directly from the vulva using a pipette or from the base of clean, empty cages. Samples were transferred to 0.5 mL Eppendorf tubes and stored at −70 °C until analysis. All urine analytes were reported as crude concentrations and as analyte-to-creatinine ratios.

#### Urine albumin to creatinine ratio

Urine samples were centrifuged at 2,000 × g for 2 minutes prior to analysis. Albumin concentrations were measured using a commercial ELISA kit (Bethyl Laboratories, Inc.). Creatinine levels were determined using a commercial assay (Roche Diagnostics) on the Cobas c 501 automated analyzer (Roche Diagnostics). The albumin-to-creatinine ratio was calculated for each sample

#### Glycated hemoglobin A1c (HbA1c)

Blood for HbA1c analysis was collected by tail vein puncture in the morning of termination. Samples were collected into heparinized glass capillary tubes and immediately suspended in Hemolyzing Reagent (Roche Diagnostics) and stored at -70°C until analysis. HbA1c was quantified using a commercial kit (Roche Diagnostics,) on the cobas c 501 autoanalyzer.

#### Analysis of circulating biomarkers of inflammation

Under isoflurane anesthesia, the abdominal cavity was surgically opened, and blood was collected from the inferior vena cava using a syringe into heparinized tubes. The samples were gently mixed by inverting the tubes five times and maintained at 4 °C until processing. Plasma was obtained by centrifugation at 3,000 × g for 10 minutes. The plasma supernatants were transferred to fresh tubes, immediately frozen on dry ice and stored at -70°C.

Plasma analytes were quantified using commercial assay kits (V-PLEX Mouse Cytokine 19-Plex Kit (Proinflammatory Panel 1 and cytokine panel 1; Catalog # K15048), Mesoscale; Mouse uPA Receptor/U-PAR ELISA kit, ab213896 Abcam). For V-PLEX Mouse Cytokine 19-Plex panel including monocyte chemoattractant protein 1 (MCP-1) and interleukin-10 (IL-10); n=13 in healthy control and n=12 in day 1, week 4, week 8 and week 12 *db/db* UNx-ReninAAV. For Cytokine Panel 1 (IL-9, MCP-1, IL-33, IL-27p28/IL-30, IL-15, IL17A/F, macrophage inflammatory protein-1 alpha (MIP-1α), interferon gamma-induced protein 10 (IP-10), and macrophage inflammatory protein-2 (MIP-2)) plasma levels were measured at a 4-fold dilution. For Proinflammatory Panel (Interferon gamma (IFN-γ), IL-1β, IL-2, IL-4, IL-5, IL-6, keratinocyte chemoattractant/growth-regulated oncogene (KC/GRO), IL-10, IL-12p70, and tumor necrosis factor alpha (TNF-α)) levels were measured at a 2-fold dilution. Values below the detection range were presented as the lower limit of quantification (LLOQ) value. For suPAR assessment, samples were randomly selected within each group; n= 9 in healthy control, n=8 in day 1 to week 8 and n=10 for week 12 *db/db* UNx-ReninAAV. Levels of suPAR were measured at a 4-fold dilution. One animal from the healthy control group was excluded as an outlier. Analysis was performed in duplicate.

### Histology

Kidney and heart were isolated at termination. The kidney capsule was removed and the whole kidney was weighed. The heart was flushed with PBS to remove excess blood before weighing. The whole heart and kidney (excluding poles) were fixed in neutral buffered formalin (NBF) containing 4% formaldehyde. After 24 hours the samples were transferred to 70% EtOH and stored at 4°C. The fixed tissue was infiltrated with paraffin overnight in a Tissue-Tek VIP6 tissue processor and embedded in paraffin blocks. Longitudinal sections of the whole heart and transverse sections of the kidney were used for histological assessment.

Glass slides with paraffin embedded sections were deparaffinated in TissueClear and rehydrated in series of graded ethanol. Immunohistochemistry (IHC) was performed using standard procedures. Briefly, after antigen retrieval and blocking of endogenous peroxidase activity, slides were incubated with primary antibody, which was subsequently detected using a polymeric HRP-linker antibody conjugate and visualized with 3,3’-Diaminobenzidine (DAB) as chromogen. Sections were counterstained in hematoxylin and cover slipped. Slides were scanned under a 40x objective in a Aperio GT450 slide scanner. Antibodies: rabbit-anti-uPA (1:100), Thermo Fischer Scientific, cat. 17968-1-AP; rabbit-anti-uPAR (heart: 1:50, kidney:1:100), Abcam, cat. ab307895; rabbit-anti-MCP-1 (1:500), Abcam, cat. ab315478; rabbit-anti-IL-10 (1:100), Thermo Fischer Scientific, cat. BS-0698R.

### Image analysis

Sections were analyzed using Visiopharm for automated quantification of immunopositive area. For renal tissues, whole kidney sections were analyzed for uPAR, uPA, and MCP-1 immunostaining. For cardiac tissues, MCP-1 and uPA immunopositive areas were quantified in longitudinal sections encompassing both the left and right ventricles. For uPAR, quantification was restricted to vascular regions with a clearly defined endothelial and smooth muscle lining in both left and right ventricle. The quantitative estimates of immunopositive staining are calculated as an area fraction (AF) in the following way:

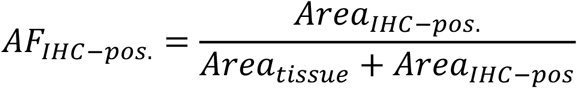

Estimation of total tissue content was calculated based on the % FA using the formula: total content (mg)= (%AF/100)*organ weight (g)*1000.

### Statistical analysis

Plotted data was analyzed using GraphPad Prism software (version 10.5.0). Data are shown as mean ± standard error of mean (SEM). For normally or lognormally distributed data, Welch’s t test (comparing two groups) or one-way ANOVA (comparing more than two groups) was applied. Dunnett’s post-hoc test was implemented for comparing means of each group to the healthy control group. For non-normally distributed data, the non-parametric statistical test Mann-Whitney (comparing two groups) or Kruskal-Wallis test (comparing more than two groups) was applied followed by a post-hoc Dunn’s multiple comparisons test. For correlation analysis a simple linear regression was used with data presented as individual values including 95% confidence bands of the best fit line. R-squared and P-values are indicated for each correlation analysis.

## Results

A total of 20 well-established circulating inflammatory cytokines and biomarkers were assessed in the *db/db* UNx-ReninAAV mouse model. The study focused on plasma sampling at timepoints when the metabolic disease phenotype is well established (32), starting at 4 weeks after UNx (defined as day 1), and thereafter analyzing timepoints 4, 8 and 12 weeks later (Supplemental Fig 1). The data was compared to *db/m* healthy control mice. Increased abundance of suPAR, MCP-1 and IL-10 were detected in the *db/db* UNx-ReninAAV mouse model in comparison to healthy control (Fig 1). Both suPAR (Fig 1C, day 1 and wk 4; P<0.0001, wk 8; P<0.001, wk 12; P<0.01) and MCP-1 (Fig 1D, day 1; P<0.05, wk 4 and 8; P<0.001, wk 12; P<0.01) were significantly elevated in the *db/db* UNx-ReninAAV model across all time-points compared to healthy control. By contrast, IL-10 levels were significantly increased only at week 12 (Fig 1E, P<0.05) relative to the healthy control group. Notably, IL-15, IL-5, IL-27p28/IL-30 and IFN-y were significantly reduced compared to healthy control in particular at earlier timepoints, from day 1 to week 4, and for IL-5 also at week 8 and for IFN-y also at week 12. The remaining 13 biomarkers were unregulated in the model (see supporting information for individual boxplots and analysis) (Fig 1, Supplemental figs 2 and 3).

**Figure 1.**
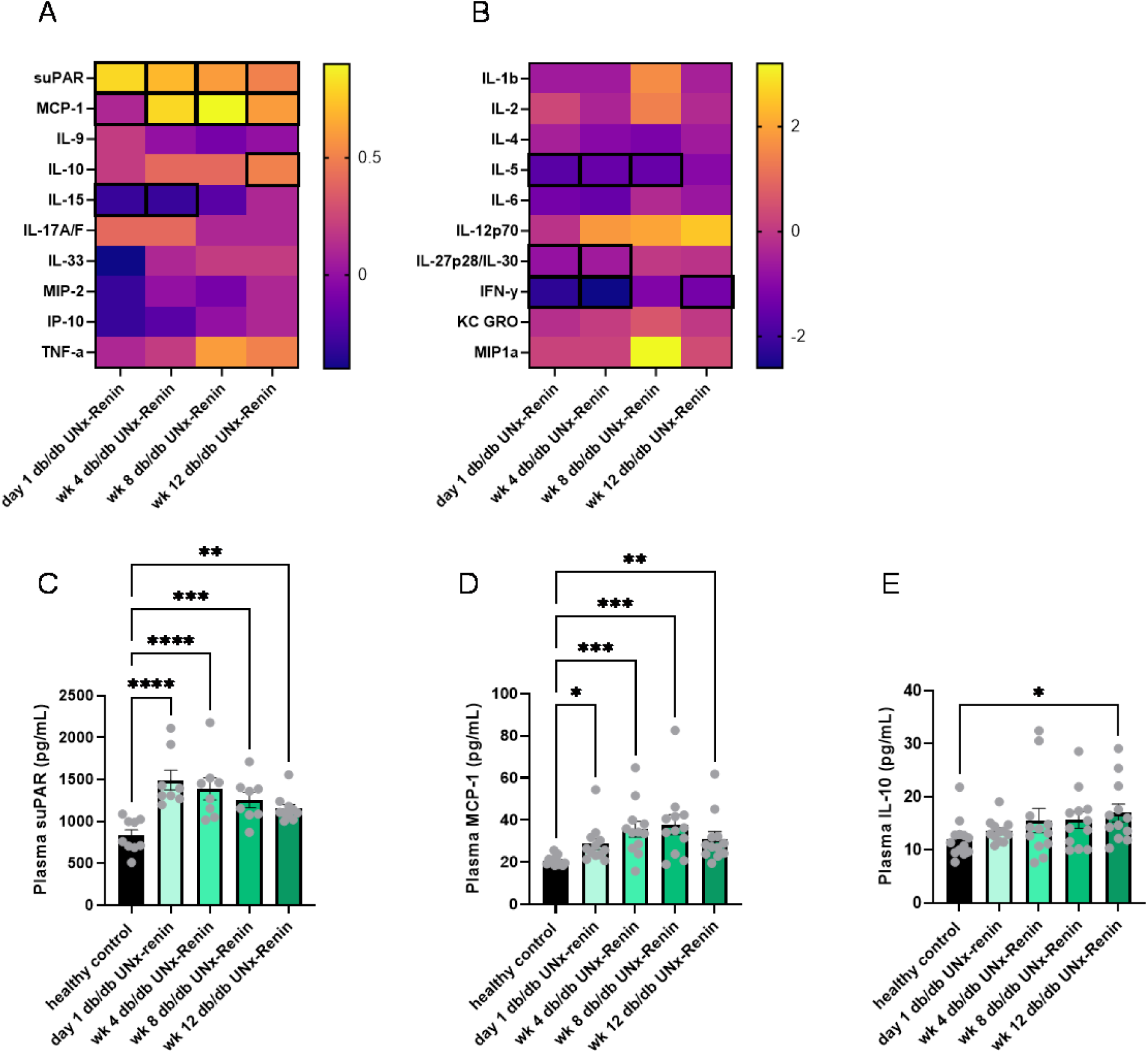
Mean log 2-fold change in circulating inflammatory biomarkers and time-course plasma levels of soluble urokinase plasminogen activator receptor (suPAR), monocyte chemoattractant protein-1 (MCP-1) and interleukin-10 (IL-10) in the *db/db* UNx-ReninAAV mouse model. Day 1 was defined as stable metabolic disease phenotype establishment, 4 weeks after UNx surgery and weeks (wk) 4, 8 and 12 are indicated in respective to Day 1. Inflammatory plasma biomarkers suPAR, MCP-1, interleukin 9 (IL-9), IL-10, IL-15, IL-17A/F, IL-33, macrophage inflammatory protein-2 (MIP-2), interferon gamma-induced protein 10 (IP-10) and tumor necrosis factor alpha (TNF-α) are presented in heatmap (A). IL-1β, IL-2, IL-4, IL-5, IL-6, IL-12p70, IL-27p28/IL-30, Interferon gamma (IFN-γ), keratinocyte chemoattractant/growth-regulated oncogene (KC GRO) and macrophage inflammatory protein-1 alpha (MIP-1α) are presented in heatmap (B). Highlighted squares (bold borders) indicate a significant difference from healthy control with P<0.05. Individual statistical analyses and boxplots are provided in Supporting information. Plasma for measuring suPAR (C), MCP-1 (D) and IL-10 (E) levels was collected at termination. n=8-13 for all data. Data is presented as means ± SEM. Lognormal ordinary one-way ANOVA with Dunnet’s post-hoc test to compare means of each group to the control group was applied to (C), Kruskal-Wallis test with Dunn’s test for multiple comparisons was used for (D-E). *p<0.05; **p<0.01***p<0.001;****p<0.0001 vs healthy control. Heat maps and graphs are generated using GraphPad Prism (v 10.5.0).

### Association of suPAR, MCP-1 and IL-10 with cardiorenal function, cardiac hypertrophy, obesity, and diabetes

To evaluate the association of inflammatory biomarkers with metabolic endpoints and cardiorenal function, plasma levels of suPAR and MCP-1 were correlated with global longitudinal strain (GLS), diastolic strain rate (reverse peak longitudinal strain rate), heart weight, body weight, uACR and HbA1c (32). Analysis was restricted to week 12 animals, when diastolic and systolic dysfunction is most evident (2,33).

suPAR levels showed significant negative correlations with GLS and diastolic strain rate (Fig 2A,B). In addition, suPAR was positively correlated with heart weight, body weight, uACR and plasma HbA1c (Fig 2C-F). Although a significant association between MCP-1 and many of the same parameters were observed using linear regression, this relationship was not robust to outlier adjustment for cardiac function endpoints (while remaining for heart weight, body weight and uACR), suggesting it may be driven by a limited number of influential observations for MCP-1 (Fig 3A-F). IL-10 levels showed weak but significant correlations with GLS, uACR, heart weight and body weight (Fig 4A-D).

**Figure 2.**
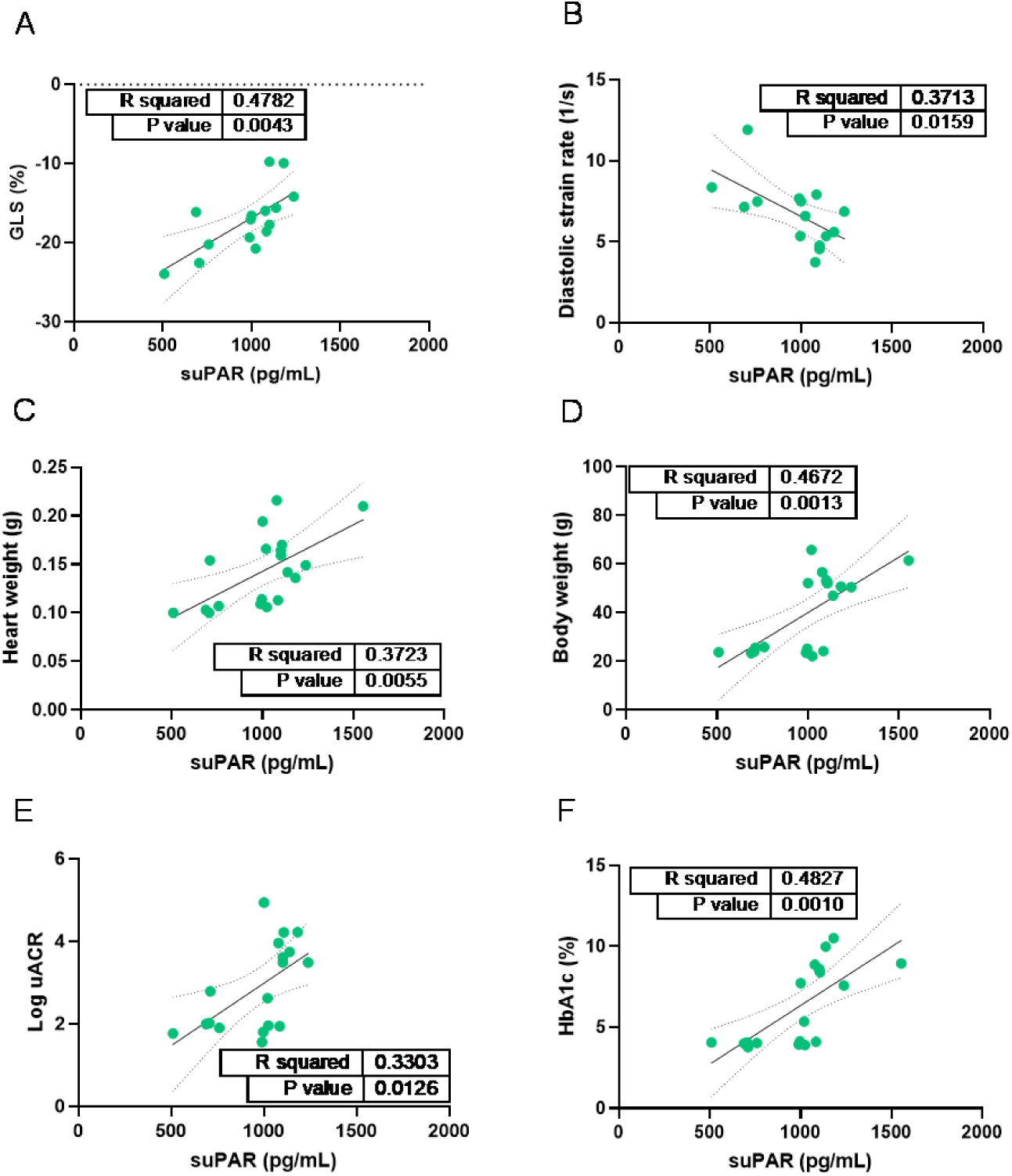
Week 12 correlation of plasma soluble urokinase plasminogen activator receptor (suPAR) with left ventricular function, cardiac hypertrophy, obesity, albuminuria and diabetes in the *db/db* UNx-ReninAAV mouse model. (A) Global longitudinal strain (GLS) and (B) diastolic strain rate were measured with echocardiography 1-2 days before termination. (C) Heart weight and (E) body weight were assessed at termination. Urine for (D) urine albumin to creatinine ratio (uACR) and blood for (F) HbA1c was obtained 1-2 days before termination and the morning of termination, respectively. n=15 for (A, B), n=18 for (C), n=19 for (D, E, F). Linear regression was used with data presented as individual values including 95% confidence bands of the best fit line. R-squared and P-values are presented for each figure. Graphs are generated using GraphPad Prism (v 10.5.0).

**Figure 3.**
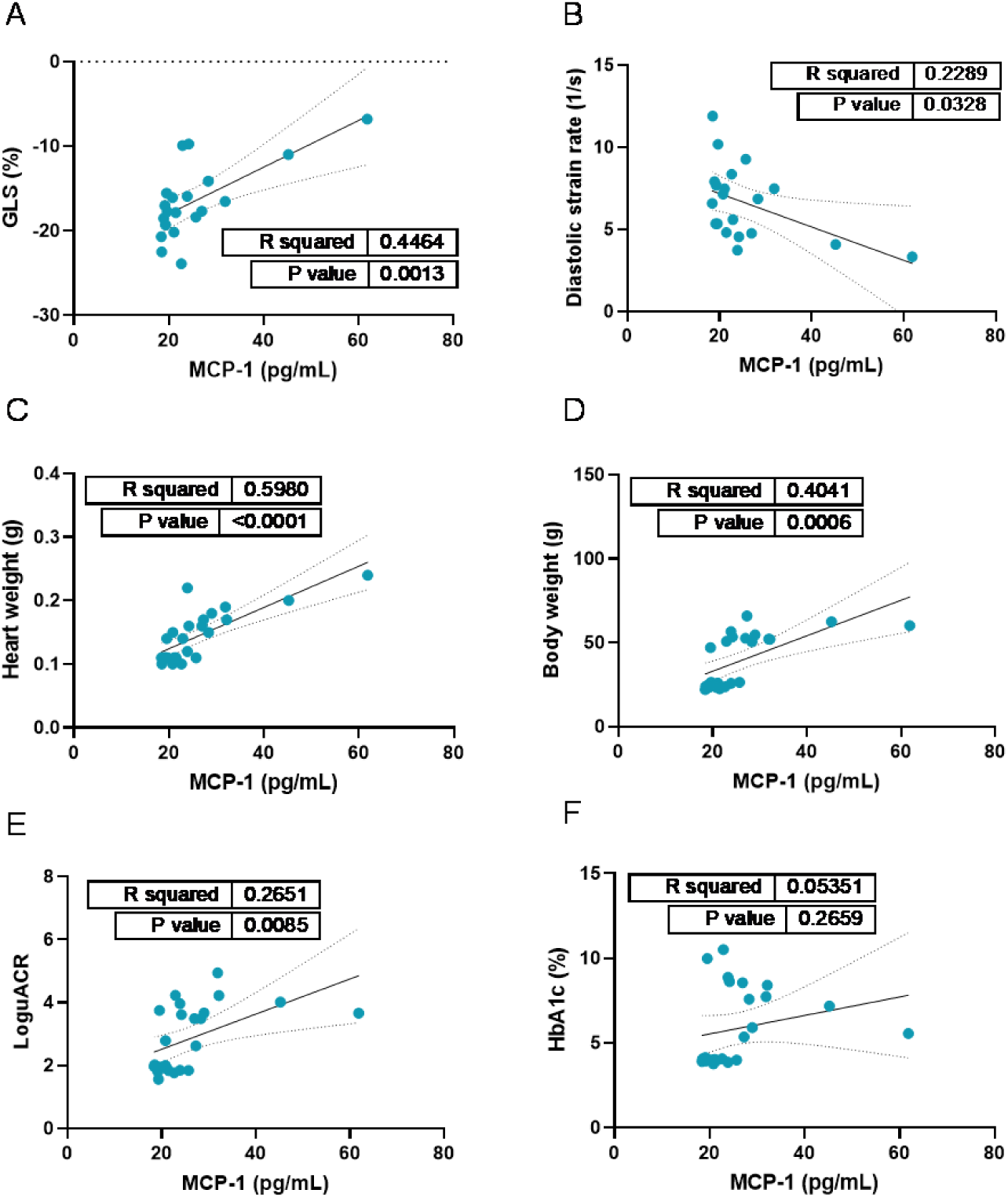
Week 12 correlation of plasma monocyte chemoattractant protein-1 (MCP-1) with left ventricular function, cardiac hypertrophy, obesity and albuminuria in the *db/db* UNx-ReninAAV mouse model. (A) Global longitudinal strain (GLS) and (B) diastolic strain rate were measured with echocardiography 1-2 days before termination. (C) Heart weight and (D) body weight were assessed at termination. Urine for (E) urine albumin to creatinine ratio (uACR) and blood for (F) HbA1c were obtained 1-2 days before termination and the morning of termination, respectively. n=20 for (A, B), n=25 for (C, D, E, F). Linear regression was used with data presented as individual values including 95% confidence bands of the best fit line. R-squared and P-values are presented for each figure. Graphs are generated using GraphPad Prism (v 10.5.0).

**Figure 4.**
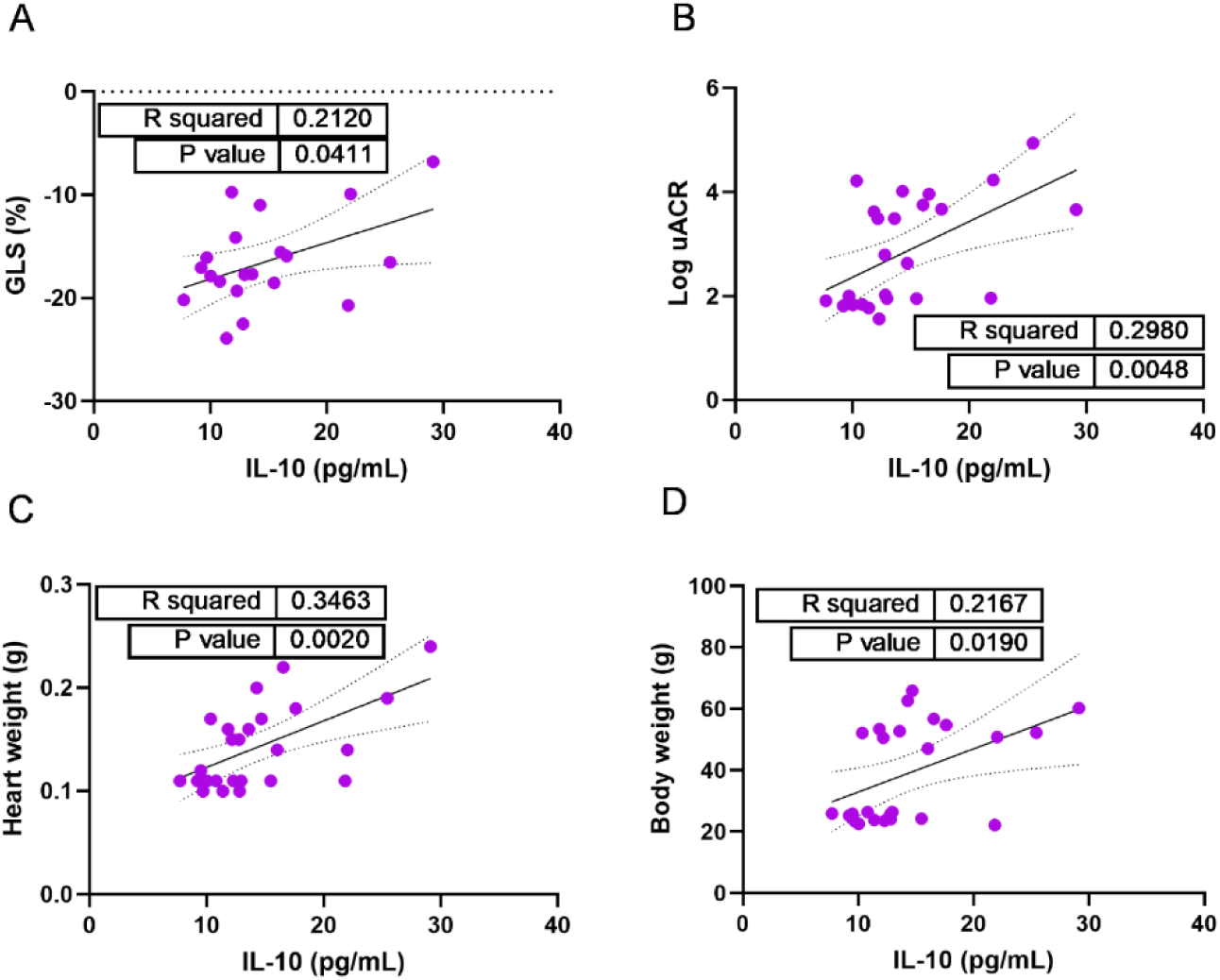
Week 12 correlation of plasma interleukin-10 (IL-10) with left ventricular function, albuminuria, cardiac hypertrophy and obesity in the *db/db* UNx-ReninAAV mouse model. (A) Global longitudinal strain (GLS) was measured with echocardiography 1-2 days before termination. Urine for (B) urine albumin to creatinine ratio (uACR) was obtained 1-2 days before termination. (C) Heart weight and (D) body weight were assessed at termination. n=20 for (A), n=25 for (B, C, D,). Linear regression was used with data presented as individual values including 95% confidence bands of the best fit line. R-squared and P-values are presented for each figure. Graphs are generated using GraphPad Prism (v 10.5.0).

### Cardiorenal expression of IL-10, uPAR, uPA and MCP-1 in the db/db UNx-ReninAAV mouse model

Given that circulating levels of IL-10, suPAR and MCP-1 were significantly elevated at week 12 in the *db/db* UNx-ReninAAV model, we next assessed whether these changes were also observed in the heart and kidney. In addition, we evaluated the expression of the uPAR ligand, urokinase plasminogen activator (uPA).

IHC staining for IL-10 revealed a diffuse distribution across both cardiac and renal tissue, suggesting either non-specific binding, expression by multiple cell types or expression outside heart and kidney (Supplemental fig 4). In contrast, uPAR expression was significantly increased in both kidney (Fig 5A-B, P<0.0001) and heart tissue (Fig 5C, P<0.01, Fig 5D, P<0.001) compared to healthy control. Representative images indicated that uPAR expression in the heart was predominantly localized to vascular wall and perivascular regions of larger vessels, whereas in the kidney it exhibited a more widespread distribution throughout the tissue, being detectable in both glomeruli and tubules (Fig 5).

**Figure 5.**
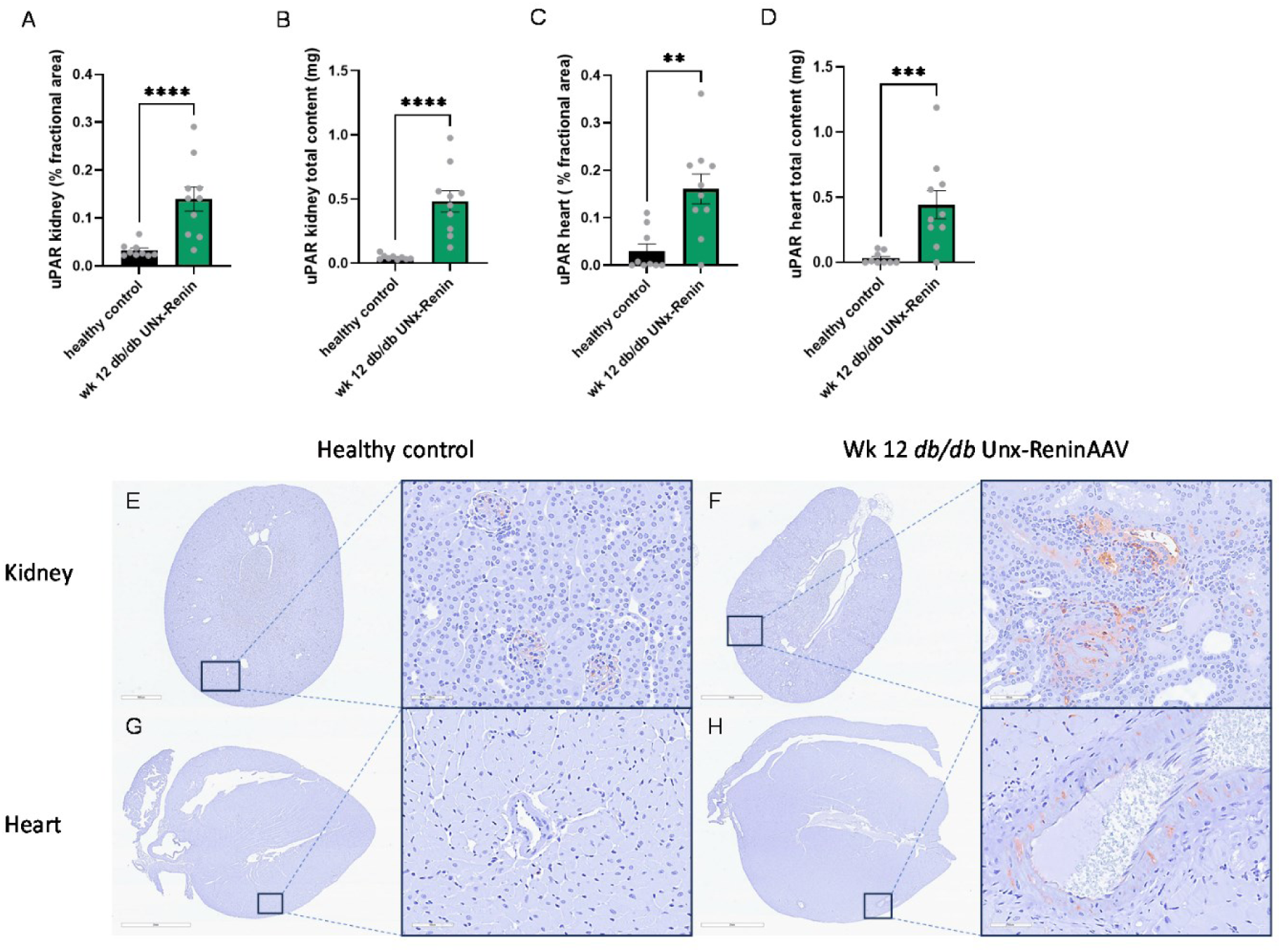
Urokinase plasminogen activator receptor (uPAR)-positive area in kidney and heart of the *db/db* UNx-ReninAAV mouse model. The uPAR positive area was assessed with immunohistochemistry (IHC) of heart and kidney sections. The percentage immunopositive area was quantified in (A) kidney and (C) heart, while total uPAR content was calculated in (B) kidney and (D) heart. N=9-10 for all data. Data is presented as means ± SEM. Lognormal Welch’s t test was applied to (A, B). Welch’s t test was applied to (C). Mann-Whitney test was used for (D). **p<0.01;***p<0.001;****p<0.0001 vs healthy control. Graphs are generated using GraphPad Prism (v 10.5.0). Representative images of the kidney and heart of healthy control (E, G) and the *db/db* UNx-ReninAAV mouse model (F, H) are presented below as whole organ view or as magnified areas (scale bar 60 µm). Negative controls are added in Supporting Information.

No significant alterations were found in the expression of the ligand uPA in heart and kidney (Fig 6A-D). Despite this, qualitative assessment of staining patterns indicated a distinct localization of uPA within the glomeruli in the kidney of *db/db* UNx-ReninAAV mice, which was not apparent in healthy controls. In the heart, uPA expression appeared to be highest in the outer myocardial layer in both model and healthy control (Fig 6).

**Figure 6.**
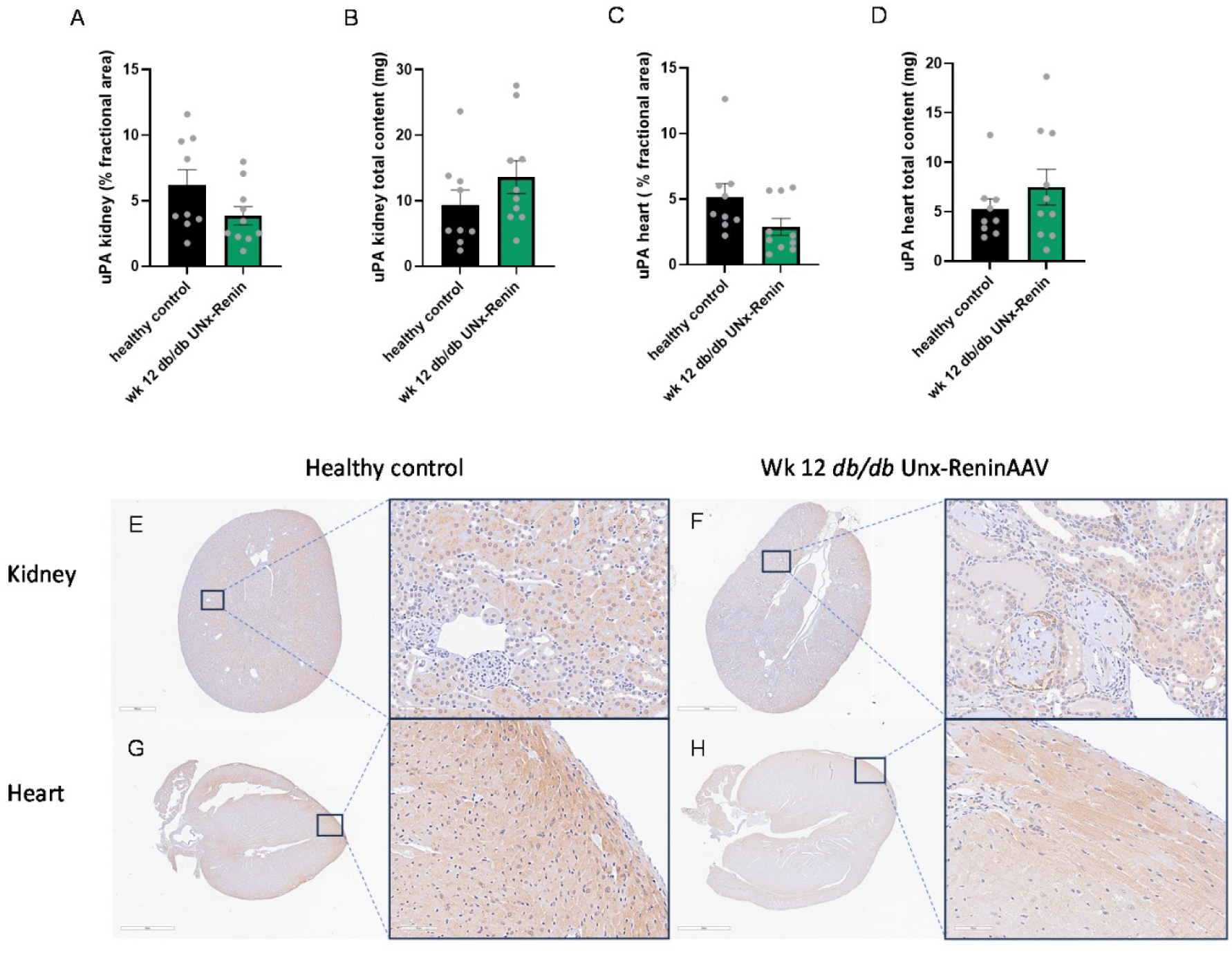
Urokinase plasminogen activator (uPA) protein expression in kidney and heart of the *db/db* UNx-ReninAAV mouse model. Protein expression of uPA was assessed with immunohistochemistry (IHC) of heart and kidney sections. % fractional area was measured in (A) kidney and (C) heart, and total content was calculated in (B) kidney and (D) heart. n=9-10 for all data. Data is presented as means ± SEM. Welch’s t test was applied to all data. Graphs are generated using GraphPad Prism (v 10.5.0). Representative images of the kidney and heart of healthy control (E, G) and the *db/db* UNx-ReninAAV mouse model (F, H) are presented below as whole organ view or as magnified areas (scale bar 70 µm). Negative controls are added in Supporting Information.

Notably, the fractional area (%) of MCP-1 expression was significantly reduced in the kidney of *db/db* UNx-ReninAAV mice compared with healthy controls (Fig 7A, P<0.0001) while the total content was significantly increased (Fig 7B P<0.01). In contrast, cardiac MCP-1 expression was markedly elevated in the model (Fig 7C-D, P<0.0001). Representative images suggested that MCP-1 expression in the kidney was predominantly localized to tubular structures, while in the heart it appeared to be more heterogeneously distributed in the vasculature and cardiomyocytes (Fig 7).

**Figure 7.**
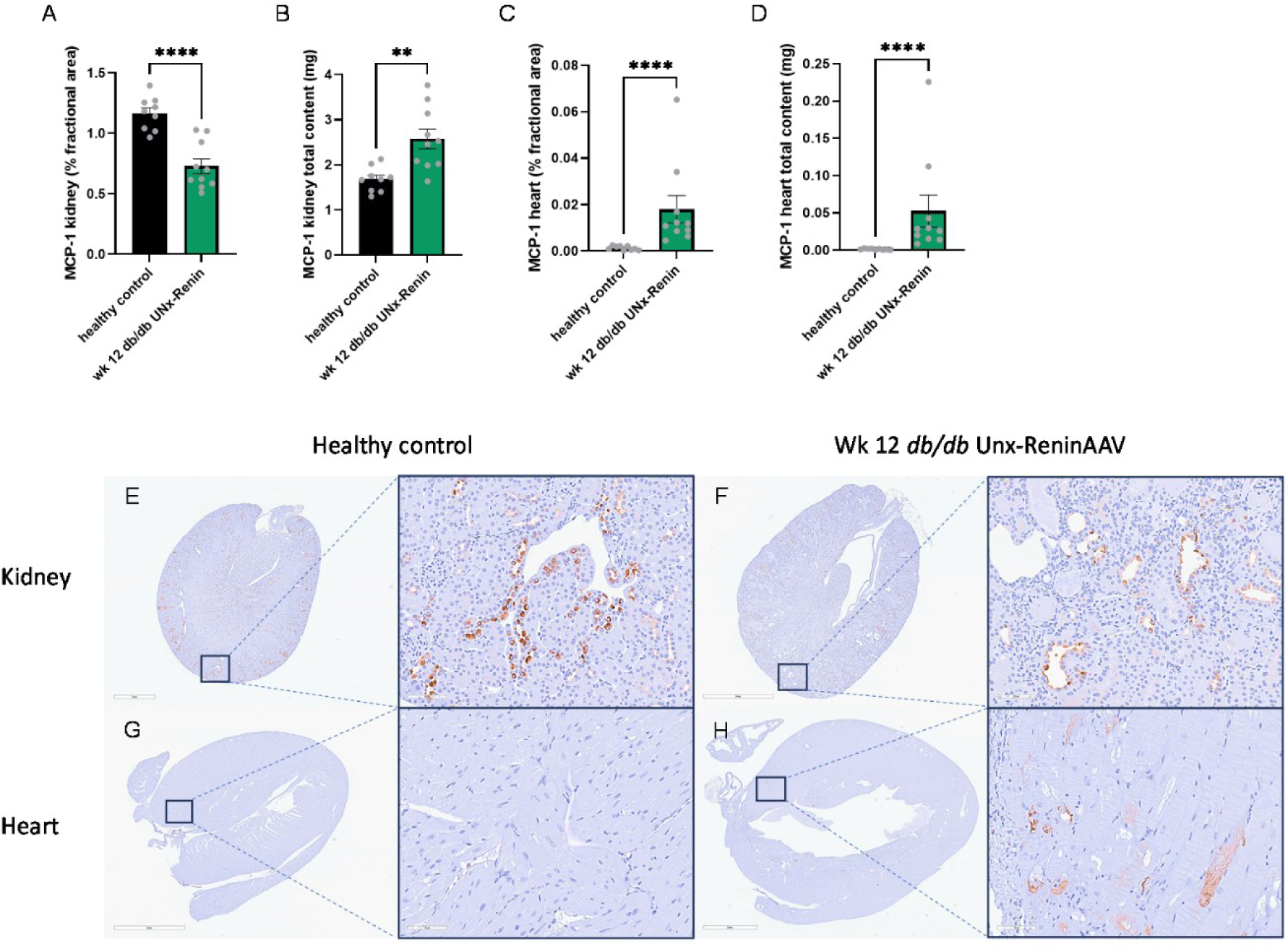
Monocyte chemoattractant protein-1 (MCP-1) protein expression in kidney and heart of the *db/db* UNx-ReninAAV mouse model. Protein expression of MCP-1 was assessed with immunohistochemistry (IHC) of heart and kidney sections. % fractional area was measured in (A) kidney and (C) heart, and total content was calculated in (B) kidney and (D) heart. n=9-10 for all data. Data is presented as means ± SEM. Welch’s t test was applied to (A). Lognormal Welch’s t test was applied to (B, D). Mann-Whitney test was used for (C). **p<0.01;****p<0.0001 vs healthy control. Graphs are generated using GraphPad Prism (v 10.5.0). Representative images of the kidney and heart of healthy control (E, G) and the *db/db* UNx-ReninAAV mouse model (F, H) are presented at the bottom as whole organ or magnified areas (scale bar 70 µm). Negative controls are added in Supporting Information.

## Discussion

Systematic analysis of translationally relevant plasma biomarkers in a mouse model combining HFpEF, CKD, and diabetes (*db/db* UNx-ReninAAV) identified a distinct inflammatory signature characterized by elevated circulating suPAR, MCP-1, and IL-10. Among these, suPAR emerged as a particularly robust marker, being consistently increased both in plasma and cardiorenal tissues. Importantly, circulating suPAR levels correlated with indices of both cardiac and renal dysfunction in the model, supporting its potential as a non-invasive, integrated biomarker of cardiorenal disease severity. MCP-1 was likewise elevated and showed associations with albuminuria, heart weight, and body weight, indicating a link to both systemic inflammation and end-organ remodeling. These findings highlight suPAR as a translationally relevant marker capable of capturing the complex pathophysiology of cardiorenal disease (33–35).

Clinically, suPAR has been extensively associated with adverse outcomes across a range of diseases (19–21,36–38). Preclinical studies have predominantly focused on the role of suPAR in kidney disease, particularly in focal segmental glomerulosclerosis (FSGS) and diabetic nephropathy, with conflicting evidence (39,40). In addition, suPAR has been associated with sepsis-induced acute kidney injury, with preclinical data indicating greater kidney damage in suPAR overexpressing mice and protective effects in suPAR deficiency (41).

In contrast, still relatively little is known about the role of suPAR in cardiovascular disease, particularly in preclinical HFpEF models. To date, mechanistic evidence in mice is largely limited to atherosclerosis, where suPAR promotes vascular inflammation, monocyte activation and plaque progression (42). Emerging evidence suggests that suPAR is not merely a biomarker but may actively contribute to cardiovascular disease through mechanisms linking chronic inflammation, endothelial dysfunction and immune activation (43). However, direct mechanistic studies in cardiac remodeling or HFpEF models remain scarce. Plasma biomarker studies in HFpEF have largely focused on natriuretic peptides, stress response markers, inflammatory cytokines and extracellular matrix remodeling associated markers (25,44,45).

At the cellular level, suPAR is the soluble form of uPAR, which is expressed by a wide range of cell types involved in immune and vascular responses. These include neutrophils, monocytes/macrophages, activated T cells, endothelial cells and fibroblasts. Circulating suPAR is generated by proteolytic cleavage of membrane-bound uPAR during inflammation and reflects the overall activity of the innate immune system (43). In the context of cardiorenal disease, both infiltrating immune cells and resident cells may contribute to increased local and systemic suPAR levels, consistent with the increased tissue staining observed in our study. Increased expression of uPAR in vascular smooth muscle cells (VSMCs) has been associated with vascular remodeling in cardiometabolic disease. uPAR has been shown to regulate VSMC phenotype by promoting the transition from a contractile to a remodeling-prone state, leading to reduced expression of contractile genes. Importantly, uPAR can exert these effects independently of its ligand uPA (46), which aligns with our findings of unchanged uPA expression in both cardiac and renal tissue. In parallel, the uPAR/suPAR system is increasingly recognized as a link between chronic inflammation, vascular dysfunction, and cardiovascular disease progression (43).

In addition to suPAR, MCP-1 and IL-10 were elevated in plasma. MCP-1, a key chemokine driving monocyte recruitment, has been widely implicated in both cardiac remodeling and progression of kidney disease (47,48). Its correlation with albuminuria, heart weight, and body weight is consistent with its known role in linking metabolic inflammation to organ damage. Interestingly, while protein expression of MCP-1 was consistently elevated in the heart, the positive area fraction was reduced in the kidney compared to healthy control. In contrast, previously investigated renal transcriptomics data from the *db/db* UNx-ReninAAV model suggests increased gene expression levels of MCP-1 (28). As the kidney tissue is severely degenerated in progressive CKD, our findings may reflect fewer viable cells in the specific area as opposed to reduced expression. IL-10, an anti-inflammatory cytokine, may reflect a compensatory response to heightened inflammatory signaling, while increased levels have been linked to macrophage induced fibroblast activation and diastolic dysfunction (49) as well as poor cardiovascular outcome and risk of CVD (50,51). However, compared to suPAR, these markers either showed fewer associations or weaker correlations with functional cardiac parameters.

Despite evidence of systemic inflammation, we did not observe elevated circulating levels of IL-6 or TNF-α, possibly reflecting disease stage-specific regulation or compensatory mechanisms. To avoid confounding effects of UNx surgery and model induction, we did not assess early time points, where some of these markers may be elevated. Interestingly, several cytokines (IL-5, IL-15, IL-27p28/IL-30, and IFN-γ) were reduced at the assessed timepoints. Reduced IL-5 and IFN-γ have been reported in chronic heart failure (52) while IL-5 and IL-15 have cardioprotective and anti-fibrotic roles (53,54). Their suppression may therefore promote fibrosis despite limited elevation of classical pro-inflammatory markers.

Together, our findings position suPAR as a promising biomarker of combined cardiorenal dysfunction in a translationally relevant model of CKD and HFpEF. Its consistent elevation across compartments, correlation with functional outcomes, and established clinical associations support further investigation into its mechanistic role and potential utility as a biomarker in cardiorenal-metabolic disease.

