## Supplemental Figures for "Circulating soluble urokinase-type plasminogen activator receptor reflects disease severity in a mouse model of diabetic kidney disease and heart failure with preserved ejection fraction"

### Supporting information

#### Supplemental figure 1

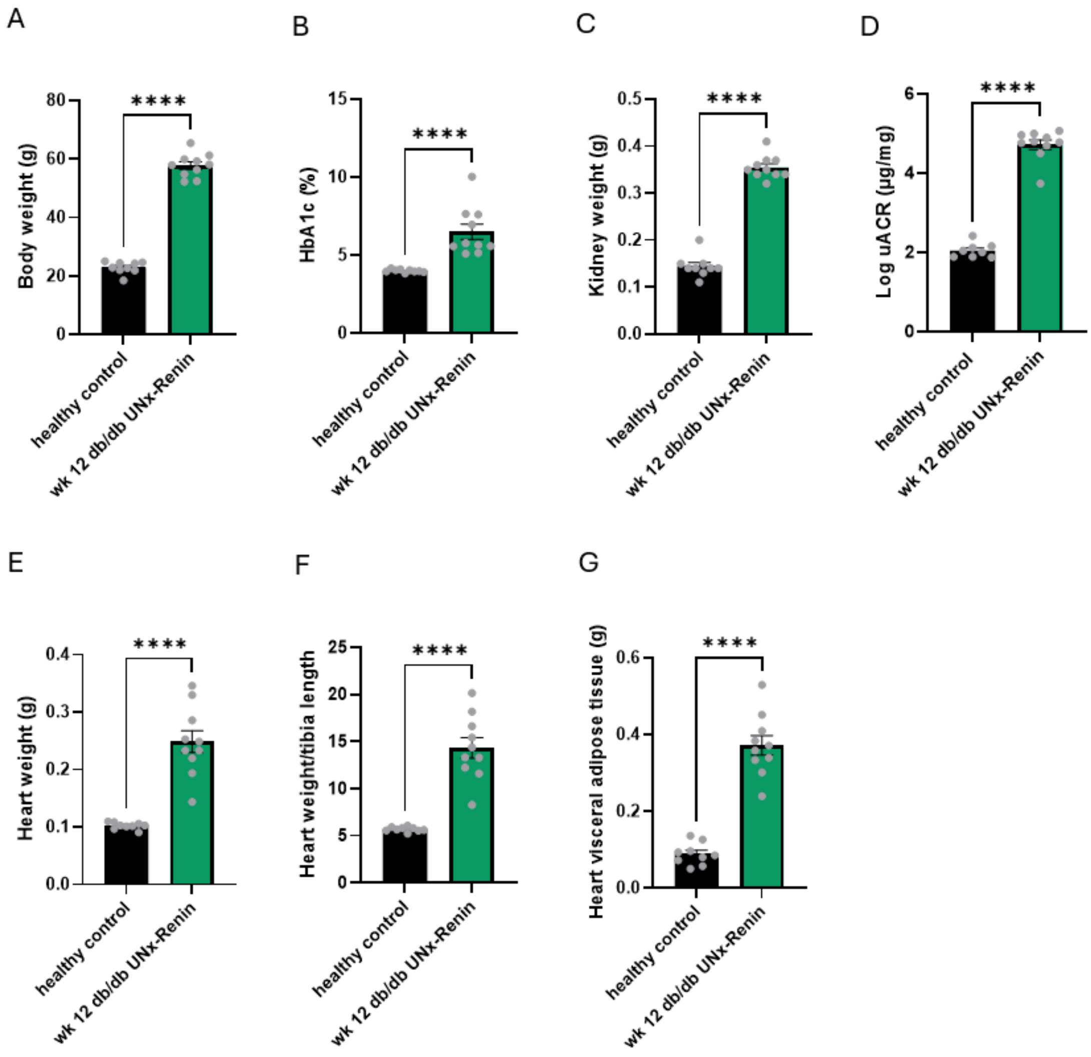

**Supplemental figure 2**

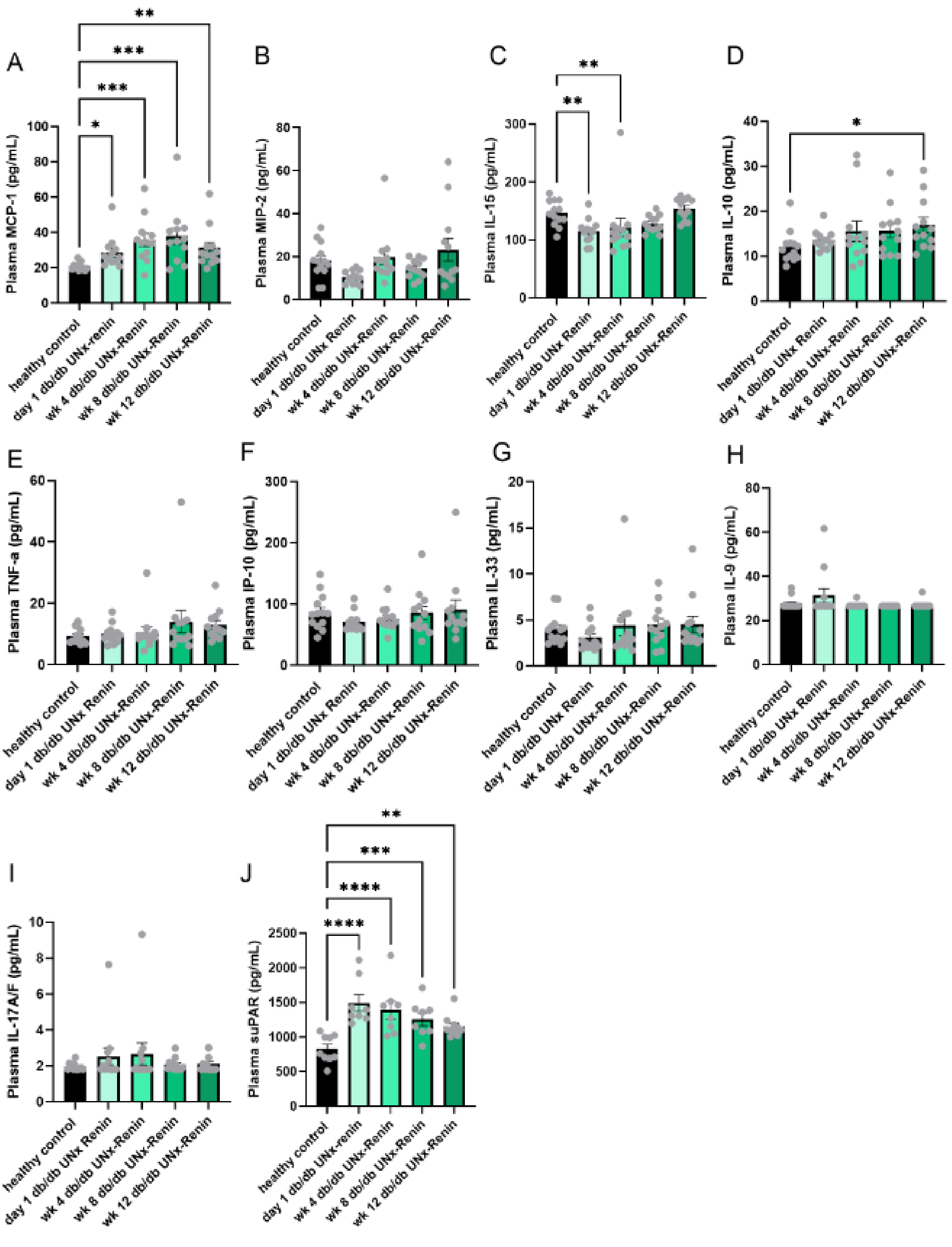

**Supplemental figure 3**

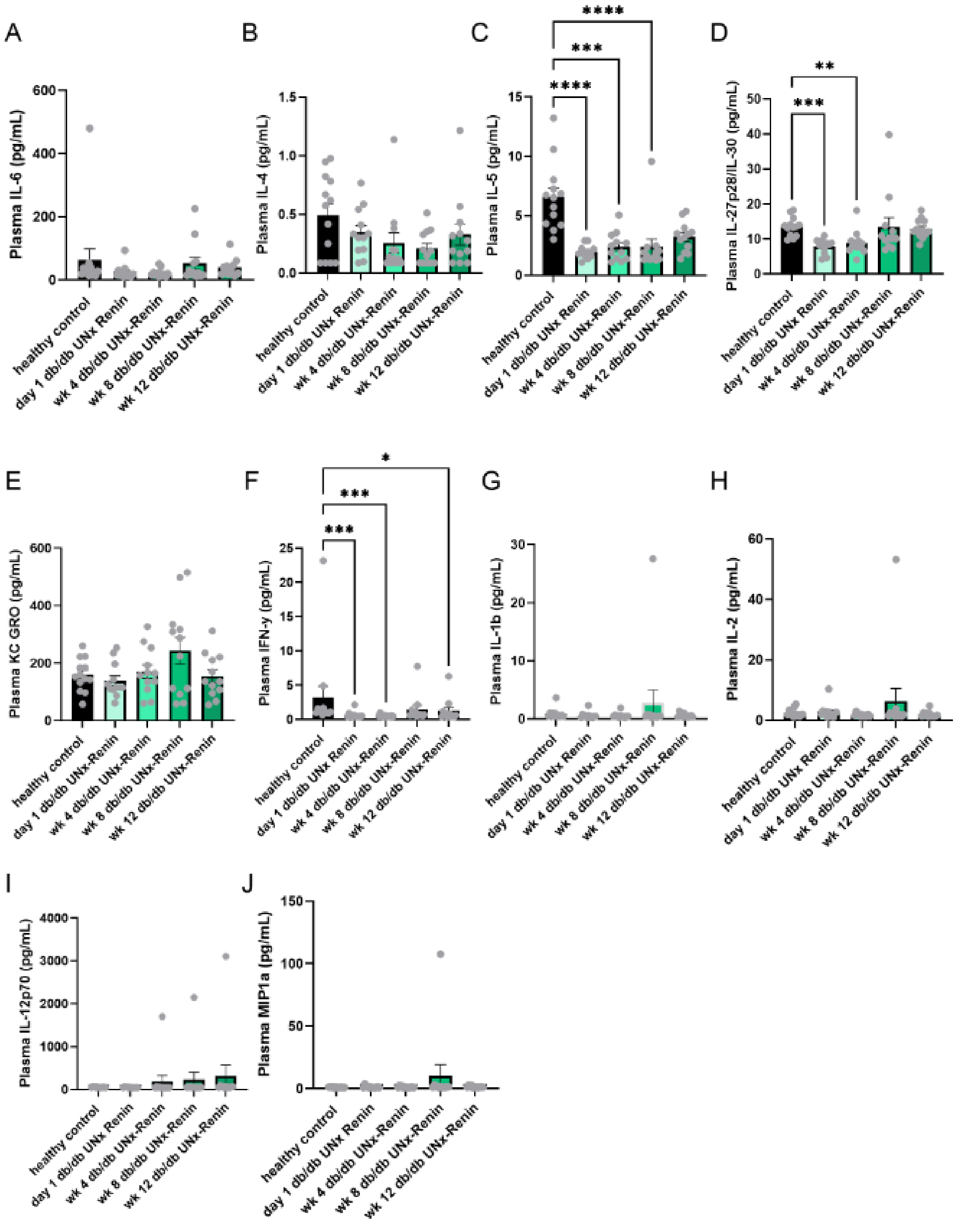

**Supplemental figure 4**

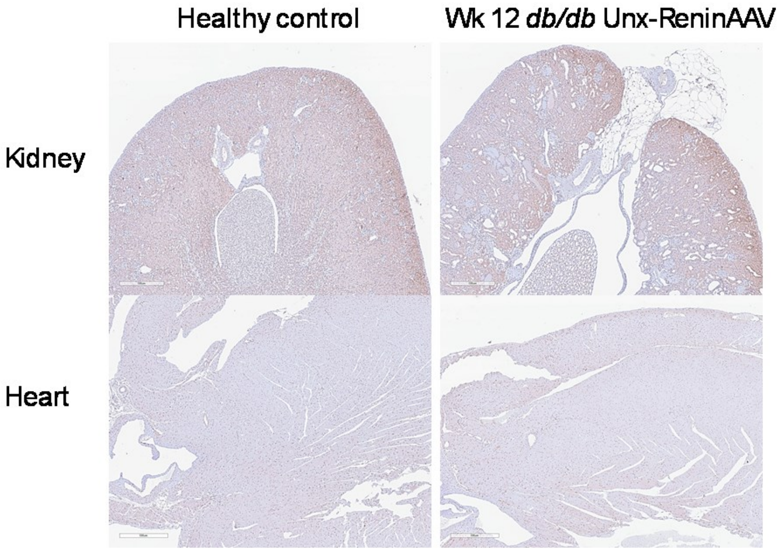

**Supplemental figure 5**

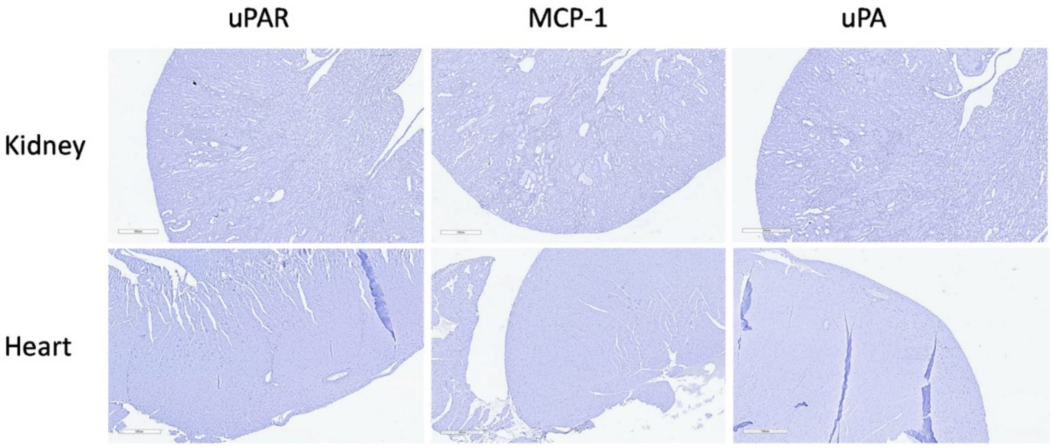

### Supplementary figure titles and legends

**Supplemental figure 1. Increased body weight, Haemoglobin A1c (HbA1c), kidney weight and urine albumin to creatinine ratio (uACR) together with cardiac hypertrophy and increased heart visceral fat in the *db/db* UNx-ReninAAV mouse model.** (A) Body weight and (C) kidney weight were measured at termination. (B) HbA1c was measured the morning of termination. (D) uACR was measured 1-2 days prior to termination. (E) Heart weight, (F) heart weight to tibia length ratio (G) and visceral fat tissue surrounding the heart were measured at termination. n=9-10 for all data. Data is presented as means ± SEM. Welch's t test was applied to (A, C, D). Mann-Whitney test was used for (B, E, F). Lognormal Welch's t test was applied to (G). **$p<0.01$;****$p<0.0001$ vs healthy control. Graphs are generated using GraphPad Prism (v 10.5.0).

**Supplemental figure 2. Circulating levels of inflammatory biomarkers in the *db/db* UNx-ReninAAV mouse model over time.** Plasma for measuring (A-J) was collected at termination and stored at -70°C until further analysis. n=8-13 for all data. Data is presented as means ± SEM. Lognormal ordinary one-way ANOVA with Dunnet's post-hoc test to compare means of each group to the control group was applied to (B, J), Kruskal-Wallis test with Dunn's test for multiple comparisons was used for (A,C-I). *$p<0.05$; **$p<0.01$***$p<0.001$;****$p<0.0001$ vs healthy control. Graphs are generated using GraphPad Prism (v 10.5.0).

**Supplemental figure 3. Circulating levels of additional inflammatory biomarkers in the *db/db* UNx-ReninAAV mouse model over time.** Plasma for measuring (A-J) was collected at termination and stored at -70°C until further analysis. n=12-13 for all data. Data is presented as means ± SEM. Lognormal ordinary one-way ANOVA with Dunnet's post-hoc test to compare means of each group to the control group was applied to (B), ordinary one-way ANOVA with Dunnet's post-hoc test to compare means of each group to the control group was applied to (E), Kruskal-Wallis test with Dunn's test for multiple comparisons was used for (A, C-J). *$p<0.05$; **$p<0.01$***$p<0.001$;****$p<0.0001$ vs healthy control. Graphs are generated using GraphPad Prism (v 10.5.0).

**Supplemental figure 4. Immunohistochemistry staining of interleukin 10 (IL-10) in kidney and heart of the *db/db* UNx-ReninAAV model.** Scale bar 500 µm.

**Supplemental figure 5. Negative controls of urokinase plasminogen activator receptor (uPAR), monocyte chemoattractant protein-1 (MCP-1) and urokinase plasminogen activator (uPA) in kidney and heart of the *db/db* UNx-ReninAAV model.** Tissue stained with second antibody only. Scale bar 500 µm.
